# ABDS: tool suite for analyzing biologically diverse samples

**DOI:** 10.1101/2023.07.05.547797

**Authors:** Dongping Du, Saurabh Bhardwaj, Sarah J. Parker, Zuolin Cheng, Zhen Zhang, Yingzhou Lu, Jennifer E. Van Eyk, Guoqiang Yu, Robert Clarke, David M. Herrington, Yue Wang

## Abstract

**Motivation:** Analytics tools are essential to identify informative molecular features about different phenotypic groups. Among the most fundamental tasks are missing value imputation, signature gene detection, and expression pattern visualization. However, most commonly used analytics tools may be problematic for characterizing biologically diverse samples when either signature genes possess uneven missing rates across different groups yet involving complex missing mechanisms, or multiple biological groups are simultaneously compared and visualized.

**Results:** We develop ABDS tool suite tailored specifically to analyzing biologically diverse samples. Mechanism-integrated group-wise imputation is developed to recruit signature genes involving informative missingness, cosine-based one-sample test is extended to detect enumerated signature genes, and unified heatmap is designed to comparably display complex expression patterns. We discuss the methodological principles and demonstrate the conceptual advantages of the three software tools. We also showcase the biomedical applications of these individual tools. Implemented in open-source R scripts, ABDS tool suite complements rather than replaces the existing tools and will allow biologists to more accurately detect interpretable molecular signals among diverse phenotypic samples.

**Availability and implementation:** The R Scripts of ABDS tool suite is freely available at https://github.com/niccolodpdu/ABDS.

**Supplementary information:** Supplementary materials are available at *Bioinformatics Advances* online.

## 1. Introduction

High-throughput molecular expression profiling technologies provide the ability to comparatively study many genes or proteins expressed in biologically diverse samples (samples belonging to different phenotypic groups) (Clarke, Ressom et al. 2008). An important but underappreciated issue in proteomics or single-cell analysis is how best to impute informative missing values that have uneven missing rates in different groups and often originate from a mix of different missing mechanisms (Shen, Chang et al. 2022). Expectedly, these missing values will have a negative impact on recruiting signature genes (Parker, Chen et al. 2020). Among many data-driven methods that have been developed to impute missing values, the categorical information associated with informative missingness is often ignored. Another essential and challenging task is to identify high quality signature genes that uniquely characterize the groups of interest against the rest. Ideally, a signature gene among molecularly distinct groups would be expressed uniquely in the individual groups of interest but in no others (Kuhn, Thu et al. 2011). However, test statistics used by most existing methods do not satisfy exactly this signature definition and are theoretically prone to detecting inaccurate signatures (Lu, Wu et al. 2022). Furthermore, while a typical heatmap design is visually effective, the common reference origin for expression measurements is altered thereafter, that is, zero-expression is replaced by floating negative values for different genes. As a result, the color coding does not rightly reflect the relative quality among signature genes.

Here we report ABDS tool suite tailored specifically to analyzing biologically diverse samples. The open-source R-script tools include mechanism-integrated group-wise missing value imputation (migImput), enumerated cosine-based one-sample test (eCOT), and unified heatmap (uniHM) design. Specifically, we propose a hybrid imputation strategy to impute informative missing values associated with signature genes, a cosine-score test on binary string references to detect enumerated signature genes, and an overall heatmap design to comparably display complex expression patterns (**Fig. 1**). We show the effectiveness and utility of these individual tools using either realistic simulations or real biomedical case studies. The ABDS tool suite will allow biologists to more accurately detect true molecular signals from biologically diverse samples.

**Figure 1.**
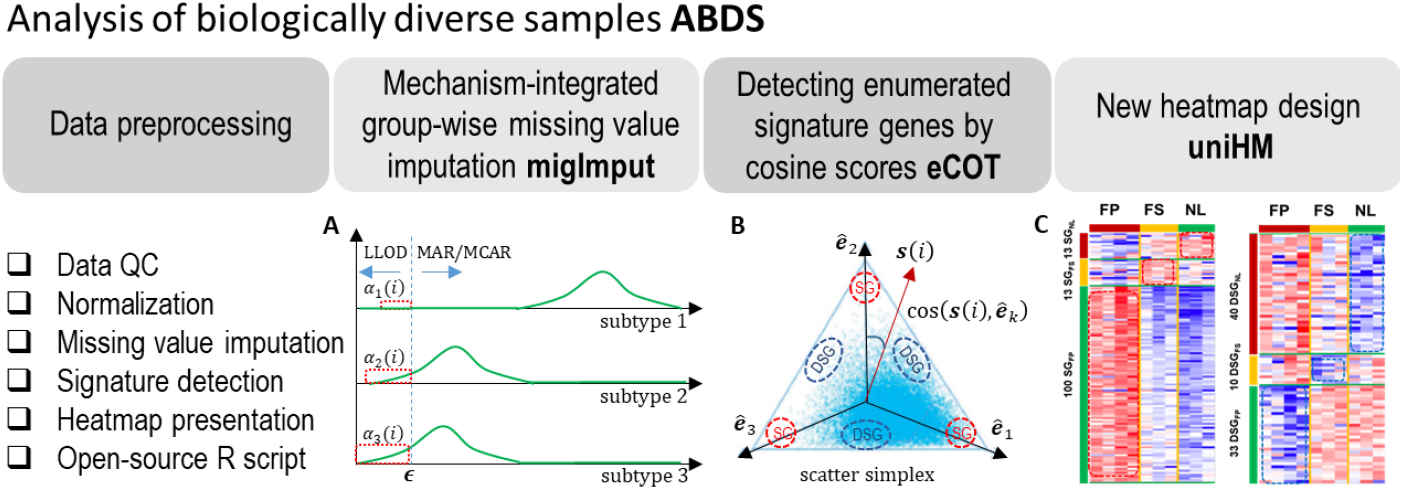
Overview of ABDS tool suite. Briefly, ABDS provides three analytics tools, namely, migImput, eCOT, and uniHM presentation.

## 2. Methods

### 2.1 Mechanism-integrated group-wise missing value imputation

Signature genes play important roles in characterizing and studying biologically diverse samples. Missing values associated with these genes are expected to have a group-specific mix of missing mechanisms and cross-group uneven missing rates. Thus, using the overall missing rate for quality control would be problematic and could adversely affect subsequent analyses including model-based missing value imputation (Shen, Chang et al. 2022). For example, missing values in the groups dominated by lower limit of detection (LLOD) are often conveniently imputed in the same way as in the groups dominated by missing at random (MAR/MCAR) mechanisms, ignoring the categorical information about biologically diverse samples (Li and Li 2018).

We propose a mechanism-integrated group-wise imputation (migImput) strategy that explicitly considers the varying mixture of missing mechanisms across different phenotypic groups. First, with an initial data normalization based on a subset of genes with no missingness, a common yet overall minimum value *ϵ* associated with LLOD is determined in log-space. Second, for each gene *i* and for each group *k*, group-specific mean value *x*_*κ*_(*i*) and standard deviation *σ*_*κ*_(*i*) are calculated. Third, within each group *k*, a missing value is imputed by (**Fig. 1A**)

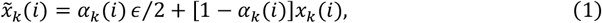

where *α*_*κ*_(*i*) is the estimated probability of the LLOD missing mechanism. The value of *α*_*κ*_(*i*) is determined by *ϵ* and the approximated normal distribution specified by *x*_*κ*_(*i*) and *σ*_*κ*_(*i*). This imputation scheme adopts ‘min/2’ for imputing LLOD missing values and ‘mean’ for imputing MAR/MCAR missing values (Shen, Chang et al. 2022) (Supplementary Information).

### 2.2 Extended cosine-based one-sample test on enumerated signature genes

Mathematically, an ideal enumerated signature gene (eSG) associated with groups *l,m,n* is defined as a gene that is expressed only in groups *l,m,n* but not in any other groups (Kuhn, Thu et al. 2011, Dai, Pei et al. 2022, Lu, Wu et al. 2022), approximately

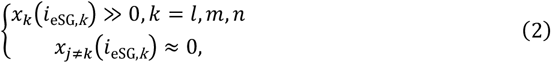

where *x*_*κ*_(*i*_eSG,*k*_) and *x*_*j*≠*κ*_(*i*_eSG,*k*_) are the average expressions of enumerated signature gene *i*_eSG,*k*_ in groups *k* and *j*, respectively. For simplicity we focus our discussion on only three groups of interest but our formulation holds for more than three groups. We emphasize that eSG as defined here are uniquely and sufficiently expressed in individual groups of interest, regardless of their actual expression level(s).

Accordingly, the cross-group expression reference of an ideal eSG can be enumerated concisely by tailored binary string ***b***_**κ**_ = [0, … ; *b*_*l*_ = 1; 0, … ; *b*_*m*_ = 1; 0, … ; *b*_*n*_ = 1; 0, …], readily for scoring *de novo* eSGs. Conceptually, the null hypothesis for non-eSG, and the alternative hypothesis for eSG, can be described as

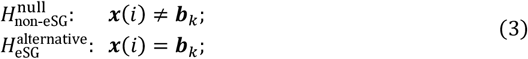

where ***x***(*i*) = [*x*_1_(*i*), *x*_2_(*i*), …, *x*_*k*_(*i*)] is the sample-averaged cross-group expression vector of gene *i*. Fundamental to the success of eCOT is the magnitude-invariant test statistic cos(***x***(*i*), ***b***_κ_) that measures directly the similarity between the cross-group expression pattern ***x***(*i*) of gene *i* and the ideal eSG expression pattern of constituent groups in scatter space (**Fig. 1B**)

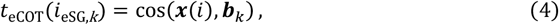

where *K* is the total number of groups. Note that two special cases of eSG are the conventional signature genes (SGs) and down-regulated signature gene (DSGs) (Kuhn, Thu et al. 2011), where SG reference is simply the Cartesian unit vectors *ê*_*κ*_ and DSG reference is 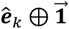 with ⊕ being the exclusive disjunction XOR operation (Supplement Information).

### 2.3 Unified heatmap design for comparative display

A popular heatmap design for displaying differentially expressed genes is to standardize each gene separately – the expression levels are first centered and then normalized by standard deviation. To address the aforementioned drawbacks, we propose an alternative heatmap design that can display both the differential pattern and referenced quality of eSGs. Specifically, for each gene *i*, the sum of group-specific mean values is calculated and used to normalize the expression level *x*(*i*) in individual samples in linear space

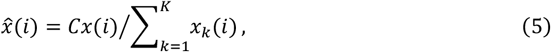

where 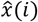 is the perspective projection of *x*(*i*) onto a scatter simplex scaled by a relatively large common constant *C*. The proximity of normalized cross-group expression vectors 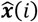 to the vertices of the scatter simplex (signature references) reflects the quality of CSGs, measured by the corresponding cosine values (Lu, Wu et al. 2022). Group-specific mean value 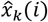 and standard deviation 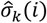 are then calculated in log-space and used to standardize the expressions of eSGs for display purpose (Supplementary Information). Furthermore, we may order sample or gene based on their sample/gene-averaged cosine values with respect to enumerated references (**Fig. 1C**) (Supplementary Information).

### 2.4 ABDS software package

The ABDS software package consists of five major analytics steps (**Fig. 1**), implemented in R scripts. The three individual tools were evaluated by community-trial software test. The R script is open-source at GitHub, and is distributed under the MIT license. The ABDS software tools are easy to use and applicable to multi-omics data. Group label on each sample is required. The output file contains the cosine scores for individual genes with respect to the ideal-pattern references of various eSGs.

## 3. Results

We conducted realistic simulation studies to evaluate the performance of three analytics tools in the ABDS package. We also conducted biomedical case studies to demonstrate the effectiveness of these tools in real-world applications.

### 3.1 migImput to recruit signature genes

We conducted simulation studies involving mixed missing mechanisms to evaluate the effectiveness of migImput tool. Our studies focused on SGs because typically these genes exhibit the highest yet most uneven missing rates or mechanisms across different groups. Group-specific SGs were generated from the truncated normal distributions centered at their ideal references (Lu, Wu et al. 2022). Missing values introduced are dominated by the random missing mechanism in the group where the SGs are highly expressed while dominated by LLOD in the groups where the SGs are lowly expressed (**Fig. 1A**). Some of the SGs were masked out according to their high overall or group-specific missing rates, and used to mimic often prematurely eliminated SGs (Supplementary Information).

We imputed missing values based on equation #1. We used both the root mean squared error (RMSE) and normalized RMSE (NRMSE) between the imputed value 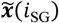 and the ground truth ***x***(*i*_SG_) to assess the imputation accuracy (Shen, Chang et al. 2022). In the comparative experiments, migImput was compared with Min/2 and Mean, the two most relevant peer methods. The experimental results are summarized in **Fig. 2** and **Table S1**. It can be seen that migImput significantly and consistently outperformed both Min/2 and Mean, in terms of both lower RMSE and NRMSE.

**Figure 2.**
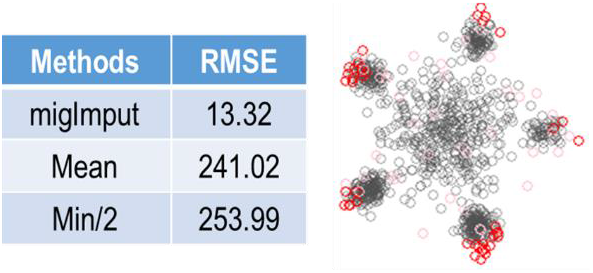
Imputation accuracy achieved by migImput as compared with Mean and Min/2 using realistic simulation studies with ground truth and measured by RMSE, where the scatter simplex shows the effective recruitment of prematurely-eliminated marker gene (color-coded) by migImput.

### 3.2 Detecting eSGs by eCOT

We conducted biomedical case studies to demonstrate the utility of eCOT tool. We first applied eCOT to a proteomics dataset acquired from human artery samples enriched by the tissue types associated with atherosclerosis (Parker, Chen et al. 2020). Samples were divided into three phenotypic groups based on the severity of atherosclerosis pathogenesis (fibrous plaque -FP, fatty streak - FS, normal - NL). We surveyed all cosine scores and reported top SGs and DSGs (**Fig. 1C, Table S2-S3**). Functional pathway analysis of tissue type-specific signature genes produced results consistent with known pathogenesis in atherogenesis. Network analysis of the top enriched functional pathways associated with FP indicated SGs were enriched for complement and coagulation functions, whereas DSGs were enriched for Myogenesis and EMT (**Fig. 3**). Together, this pattern is consistent with increased inflammation and decreased smooth muscle cell contractile phenotype composition within atherosclerotic lesions. Pathway analysis also indicated association with mTORC1 signaling and reactive oxygen species pathway enriched in FS and myogenesis, EMT, hypoxia and IL2/STAT5 signaling in NL (**Fig. S1**), all previously been linked to atherogenesis. Since IL2-related DSGs were enriched in the NL, this finding could reflect that lower IL2 signaling is protective against atherosclerotic plaque development.

**Figure 3.**
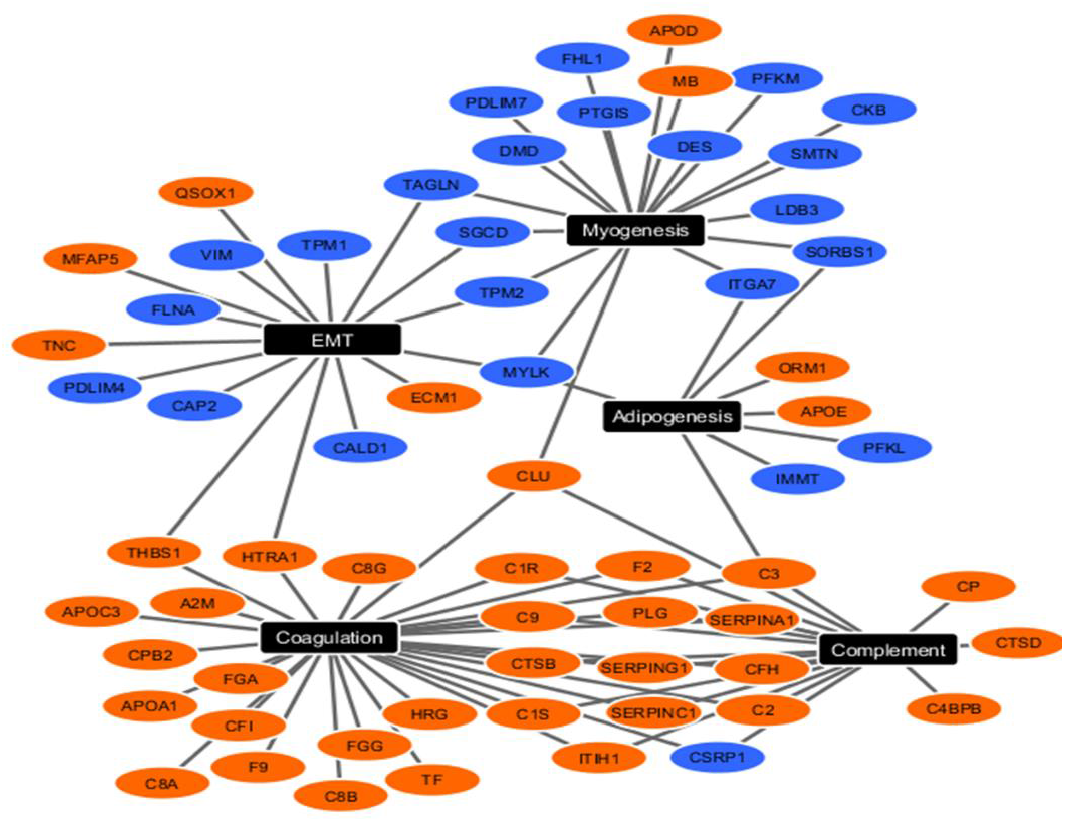
Upregulated (orange nodes) and downregulated (blue nodes) SGs/DSGs in FP group clustered into the top 5 functional pathways from the MSigDB component of Enrichr pathway analysis software.

We then applied eCOT to the Edinburgh breast cancer gene expression data that were acquired prior to standard treatment. Samples were divided into four roughly equal-sized phenotypic groups based on the follow-up sample-wise recurrence times (Supplementary Information). We again surveyed all possible forms of eSGs and reported top eSGs (**Figs. S2-S4, Tables S4-S6**). Signaling activated downstream of EGFR family members is a central feature of breast cancer. HER2/ERBB2 is the most widely studied, where protein overexpression or gene amplification defines one of the three primary breast cancer groups and targeting the HER2 protein and/or blocking its kinase function greatly improves overall survival is now standard of care for patients with HER2+ breast tumors. When applied to transcriptome data from breast cancer patients, the eCOT identified EGFR/ERBB2 and multiple EGFR-related downstream targets as enriched in estrogen-receptor positive (ER+) breast cancers likely to recur late (≥ 5 years after initial diagnosis). Most of these tumors were treated with the antiestrogen Tamoxifen and many patients would experience an overall survival benefit from Tamoxifen. However, consistent with the eCOT prediction, higher expression of EGFR (ERBB) or HER2 (ERBB2) would be expected to reduce Tamoxifen responsiveness and increase the likelihood of a subsequent recurrence.

### 3.3 Visualizing eSGs by uniHM

We used the newly designed heatmap to display the differential expression patterns of the eSGs reported in section 3.2 (**Fig. 1C, Figs. S2-S4**), in comparison with the classically designed heatmap (**Fig. S5**). Using this newly implemented heatmap function, eSGs are arranged based on their sample-averaged cosine scores with respect to hypothesis-enumerated references. The new heatmap visually reflects the idealness of eSGs where the common origin remains the same across all genes and the contrast is consistent with the corresponding cosine scoring.

## 4. Discussion

The ABDS tool suite provides three data analytics tools tailored for analyzing biologically diverse samples across many groups. The newly introduced tools complement the existing tools for imputing mechanism-mixed informative missing values, detecting group-enumerated signature genes, and visualizing complex differential expression patterns. Specifically, migImput will help recruit critical SGs that are often prematurely eliminated due to high overall missing rates. Moreover, the detected eSGs will allow scientists to test more and specific hypotheses, potentially providing an informative subspace for disease progression mapping. Note that eSGs are different from regrouped differentially expressed genes (Kuhn, Thu et al. 2011). For readers interested in the algorithmic workflows and comparative evaluations of these tools versus peer methods, we highly recommend the relevant reports (Dai, Pei et al. 2022, Shen, Chang et al. 2022).

We emphasize that the ABDS tool suite complements rather than replaces existing tools. For example, within each group, migImput imputes potentially informative missing values by considering both LLOD and MAR/MCAR mechanisms and our limited comparisons have focused on the two most relevant peer methods (Min/2 and Mean). We note that assessing imputation accuracy over masked values is intrinsically limited for real data because evaluation is not directly over authentic missing values (they will never be known). Hence, we recommend that users may apply additional global methods to refine missing value imputation after migImput (Li and Li 2018, Shen, Chang et al. 2022). We also advise users to apply a classical heatmap design for visualizing differential expression patterns.

**Table 1.** Imputation accuracy achieved by migImput as compared with Mean and Min/2 using realistic simulation studies with ground truth and measured by NRMSE. In this experiment, the missing rate threshold for masking-out is 60% (overall), resulting in a total of 83 features masked-out, including 44 SGs and 39 non-SGs, containing 4,355 missing values

|  | migImput | Mean | Min/2 |
| --- | --- | --- | --- |
| NRMSE | 0.0406 | 0.7348 | 0.7743 |

## ACKNOWLEDGMENTS/FUNDING

This work has been supported by the National Institutes of Health under Grants HL111362, HL133932, CA271891, MH110504, NS123719, and the Department of Defence under Grant BC171885P1.

## COMPETING FINANCIAL INTERESTS

The authors declare no competing financial interests.

## Data availability

Human artery proteomics dataset was obtained from publicly available datasets from previously published study available at https://pubs.acs.org/doi/10.1021/acs.jproteome.0c00118 (Parker, Chen et al. 2020).

## Notes

https://github.com/niccolodpdu/ABDS

### Competing Interest Statement

The authors have declared no competing interest.

## REFERENCES

Clarke, R., H. W. Ressom, A. Wang, J. Xuan, M. C. Liu, E. A. Gehan and Y. Wang (2008). “The properties of high-dimensional data spaces: implications for exploring gene and protein expression data.” Nat Rev Cancer 8(1): 37–49.

Dai, M., X. Pei and X. J. Wang (2022). “Accurate and fast cell marker gene identification with COSG.” Brief Bioinform 23(2).

Kuhn, A., D. Thu, H. J. Waldvogel, R. L. Faull and R. Luthi-Carter (2011). “Population-specific expression analysis (PSEA) reveals molecular changes in diseased brain.” Nat Methods 8(11): 945–947.

Li, W. V. and J. J. Li (2018). “An accurate and robust imputation method scImpute for single-cell RNA-seq data.” Nat Commun 9(1): 997.

Lu, Y., C. T. Wu, S. J. Parker, Z. Cheng, G. Saylor, J. E. Van Eyk, G. Yu, R. Clarke, D. M. Herrington and Y. Wang (2022). “COT: an efficient and accurate method for detecting marker genes among many subtypes.” Bioinform Adv 2(1): vbac037.

Parker, S. J., L. Chen, W. Spivia, G. Saylor, C. Mao, V. Venkatraman, R. J. Holewinski, M. Mastali, R. Pandey, G. Athas, G. Yu, Q. Fu, D. Troxlair, R. Vander Heide, D. Herrington, J. E. Van Eyk and Y. Wang (2020). “Identification of putative early atherosclerosis biomarkers by unsupervised deconvolution of heterogeneous vascular proteomes.” J Proteome Res 19(7): 2794–2806.

Shen, M., Y. T. Chang, C. T. Wu, S. J. Parker, G. Saylor, Y. Wang, G. Yu, J. E. Van Eyk, R. Clarke, D. M. Herrington and Y. Wang (2022). “Comparative assessment and novel strategy on methods for imputing proteomics data.” Sci Rep 12(1): 1067.

