## Supplementary Information for "ABDS: tool suite for analyzing biologically diverse samples"

### Methods

#### Background and Motivation

High-throughput molecular expression profiling technologies provide the ability to comparatively study many genes or proteins expressed in biologically diverse samples (samples belonging to different phenotypic groups) (Clarke, et al., 2008). Among the most fundamental tasks are missing value imputation, signature gene detection, and expression pattern visualization. However, most commonly used analytics tools may be problematic for characterizing biologically diverse samples when either signature genes possess uneven missing rates across different groups yet involving complex missing mechanisms, or multiple biological groups are simultaneously compared and visualized.

While there are multiple causes for missing values in omics data, three typical missing mechanisms are widely acknowledged. For example, low abundant proteins may be missing because their concentration is below the lower limit of detection (LLOD); while poorly ionizing peptides or problems in technical pre-processing may cause proteins to be missing not at random (MNAR) (Herrington, et al., 2018). However, missingness may also extend to mid- and even high-range intensities (Dabke, et al., 2021), statistically categorized into missing at random (MAR) and missing completely at random (MCAR) (Lazar, et al., 2016). MAR is actually missing conditionally at random given the observed data distribution or underlying parametric covariates. MCAR depends on neither observed nor missing data, thus the incomplete data are representative of the entire dataset. While MAR allows prediction of the missing values based on observed data, unfortunately, the MAR and MNAR conditions cannot be distinguished based on the observed data because by definition missing values are unknown (Jakobsen, et al., 2017; Liu and Dongre, 2020). More importantly, missing values in reality can originate from a mix of both known and unknown missing mechanisms (Lazar, et al., 2016; Webb-Robertson, et al., 2015). A common solution for missingness is to impute the missing values based on assumed missing mechanisms. However, this approach can introduce a profound change in the distribution of protein-level intensities because most methods are only designed for a single missing mechanism. These changes can have unpredictable effects on downstream differential analyses.

To explore a more integrative strategy for improving imputation performance, we discuss a low-rank matrix factorization framework with fused regularization on both sparsity and similarity – Fused Regularization Matrix factorization (Lin and Boutros, 2020; Ma, et al., 2011; Wang, et al., 2016), which can naturally integrate other-omics data such as gene expression or clinical variables. We also introduce a biologically-inspired latent variable modeling strategy - Convex Analysis of Mixtures (Chen, et al., 2020; Wang, et al., 2016), which explicitly formulates a data matrix as mixtures of underlying biological archetypes and performs missing value imputation on original intensity data (before log-transformation). Preliminary results on real proteomics data are provided together with an outlook into future development directions.

An important but frequently underappreciated issue is how best to define and detect a cell or tissue marker among many groups. Ideally, a molecularly distinct group would be composed of molecular features that are expressed uniquely in the cell or tissue group of interest but in no others – so called marker genes (MG) (Kuhn, et al., 2011). With the increasing availability of group expression profiles acquired by single or sorted cell sequencing, data-driven software tools to detect MG are essential for characterization, classification, or deconvolution of tissue or cell groups (Herrington, et al., 2018; Parker, et al., 2020).

The most frequently used methods rely on an ANOVA model that adopts the null hypothesis that samples in all groups are drawn from the same population and is originally designed to detect differentially-expressed genes across any of the groups. Another popular method is the One-Versus-Rest Fold Change or t-test (OVR-FC/t-test/Limma/EdgeR) that is based on the ratio of the averaged expression in a particular group to the averaged expression in all other groups (Chikina, et al., 2015; Ritchie, et al., 2015). However, a gene with a low average expression value in the rest is not necessarily expressed at a low level in every group in the rest. An alternative strategy is the One-Versus-Everyone Fold Change (OVE-FC) or its variants (Chen, et al., 2021; Newman, et al., 2015). Because an OVE test compares only the top two groups (with the highest or second-highest averaged expression value) for mathematical convenience, the remaining informative groups are neglected. We and others have recognized that these test statistics used by most current methods do not satisfy exactly the MG definition and are theoretically prone to detecting inaccurate MG (Kuhn, et al., 2011).

We previously reported an accurate and efficient data-driven method - Cosine based One-sample Test (COT) - to detect MG among many groups (Lu, et al., 2022). Note that eCOT is an extended version of COT. Formulated as a one-sample test, the test statistic of COT is the cosine similarity between a molecule’s expression pattern across all groups and the exact mathematical definition of an ideal MG. Under the assumption that most genes are associated with the null hypothesis, COT approximates the empirical null distribution with a finite normal mixture distribution for calculating p-values (Efron, 2004). We implemented the COT workflow in a Python package, evaluated and compared MG detection by COT and peer methods using realistic simulation data (Lu, et al., 2022).

**Mechanism-integrated group-wise imputation by migImput**

The current practice in analyzing omics data containing missing values is to eliminate genes with overall missing rates higher than a threshold. This would be problematic when samples are biologically diverse, e.g. belonging to multiple yet different groups. The proposed migImput aims to pre-impute the missing values particularly associated with marker or signature genes across groups. Thus, it complements rather than replace existing methods. The key difference between migImput and existing methods is that migImput performs group-wise imputation, integrates both MNAR and MAR/MCAR mechanisms, and aims to retain marker or signatures genes that may be eliminated prematurely due to relatively high overall missing rates (so-called informative missingness). The most logical or closely relevant peer methods would be Min/2 and Mean (Shen, et al., 2022). Our objective evaluation and limited comparison show the effectiveness of migImput for achieving the designed objectives, in particular, for retaining marker or signature genes that are critical in characterizing or deconvoluting bulk samples of mixed groups.

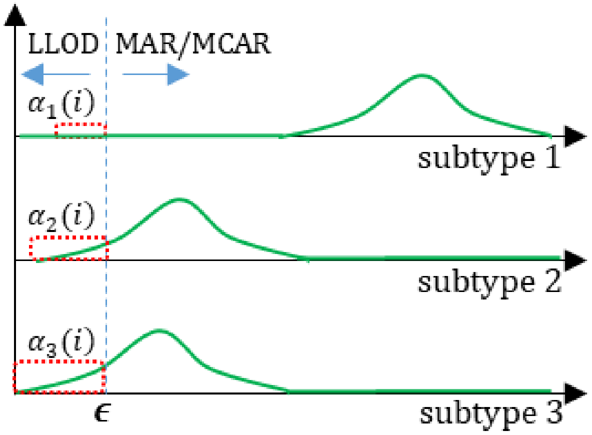

Conceptually, Min/2 can be used to impute missing values due to LLOD (MNAR) mechanism, and Mean can be used to impute missing values due to MAR/MCAR. To pre-impute marker or signature genes associated with different groups with uneven missing rate or mechanism-mix (see above illustration), for each group k, we plug-in the global minimum to the estimated normal distribution to calculate the probability $\alpha_k(i)$ of LLOD/MNAR (red area), then assign $[1-\alpha_k(i)]$ to be the probability of MCAR/MAR. We use group-specific mean and global Min/2 for group-specific imputation based on equation #1 in the main text.

In relation to the Mean imputation strategy, the important thing is that global mean may bias groups differences toward the null, and this bias may be more pronounced if there is asymmetry between the number of samples in each group. The same effect can occur if group differences vary by batch. Consider a toy example: 50% missingness, observed mean values FP=4, FS=2, NL=2; the global mean is 8/3; then the imputed mean values FP=3.33, FS=2.33, NL=2.33. For another gene (more differential), the observed mean values FP=4, FS=1, NL=1; the global mean is 2; the imputed mean values FP=3, FS=2, NL=2. In contrast, instead of using global mean, migImput uses group-specific mean (together with Min/2) to impute group-specific missing values. In the toy example, group-specific means imputed by migImput remain unchanged.

**Original concept of COT for detecting signature genes**

As aforementioned, eCOT is an extended version COT. Here we introduce and discuss the original concept of COT. Mathematically, an ideal MG of group *k* is defined as a gene expressed only in group *k* but not in any other groups (Chikina, et al., 2015; Delaney, et al., 2019; Kuhn, et al., 2011; Lu, et al., 2022), approximately

$$\begin{cases} s_k(i_{\mathrm{MG},k}) \gg 0, \\ s_{l\neq k}(i_{\mathrm{MG},k}) \approx 0, \end{cases} \tag{1}$$

where $s_k(i_{\mathrm{MG},k})$ and $s_{l\neq k}(i_{\mathrm{MG},k})$ are the average expressions of marker gene $i_{\mathrm{MG},k}$ in groups *k* and $l$, respectively. We acknowledge that there are alternative definitions but in the absence of a universally accepted standard in the field, our definition provides some unique advantages to guide our work. We emphasize that group-specific MG as defined here are enriched uniquely in a particular group, regardless of their expression level(s), and their identities can be readily used in facilitating deconvolution or classification (Wang, et al., 2016).

Accordingly, the cross-group expression pattern of an ideal MG can be represented concisely by the Cartesian unit vectors $\hat{\boldsymbol{e}}_k$, readily serving as a reference for a one-sample test.

Conceptually, the null hypothesis for non-MG, and the alternative hypothesis for MG, can be described as

$$\begin{aligned} H_{\text{non-MG}}^{\text{null}}&: \quad \boldsymbol{s}(i) \neq \hat{\boldsymbol{e}}_k; \\ H_{\text{MG}}^{\text{alternative}}&: \quad \boldsymbol{s}(i) = \hat{\boldsymbol{e}}_k; \end{aligned} \tag{2}$$

where $\boldsymbol{s}(i) = [s_1(i), s_2(i), \dots, s_K(i)]$ is the sample-averaged cross-group expression pattern of gene $i$. Fundamental to the success of COT is the newly-proposed test statistic $\cos(\boldsymbol{s}(i), \hat{\boldsymbol{e}}_k)$ that measures directly the similarity between the cross-group expression pattern $\boldsymbol{s}(i)$ of gene $i$ and the ideal MG expression pattern of constituent groups in scatter space given by

$$t_{\text{COT}}(i_{\text{MG}}) = \underset{1 \le k \le K}{\operatorname{argmax}} \cos(\boldsymbol{s}(i), \hat{\boldsymbol{e}}_k) = \underset{1 \le k \le K}{\operatorname{argmax}} \frac{s_k(i)}{\sqrt{\sum_{j=1}^{K} [s_j(i)]^2}}, \tag{3}$$

where $K$ is the number of constituent groups. Sample normalization and batch effect adjustment are the required preprocessing steps prior to COT analysis; when applicable, the input of COT should be a sample-normalized and batch-adjusted data matrix.

We implemented the COT workflow in both Python and R, and used community-based trials to test the COT software. The Python package is open-source at GitHub, built using NumPy and Pandas, and is distributed under the MIT license. The COT software tool is easy to use and applicable to multi-omics data. The rows of input data matrix correspond to genes or other molecular features, and the columns correspond to samples. The group label on each sample is required by the COT test statistic. The output file stores the input genes and their cosine values in reference to the ideal MG of respective groups (Lu, et al., 2022).

**Performance index**

Two quantitative measures were used to evaluate imputation accuracy, namely Root Mean Square Error (RMSE) and Normalized Root Mean Square Error (NRMSE). Specifically, RMSE and NRMSE are given by (Oba, et al., 2003; Stekhoven and Bühlmann, 2012)

$$\text{RMSE} = \sqrt{\frac{\sum_{\Omega}(\hat{X}_{\Omega} - X_{\Omega})^2}{|\Omega|}}, \quad \text{NRMSE} = \sqrt{\frac{\sum_{\Omega}(\hat{X}_{\Omega} - X_{\Omega})^2}{|\Omega| \sigma_{X_{\Omega}}^2}},$$

respectively, where $\Omega$ is the index set of missing values in complete data matrix $X$, $|\Omega|$ is the total number of missing values, $\hat{X}$ is the imputed complete data matrix, and $\sigma_{X_{\Omega}}^2$ is the variance of missing values.

##### Two most-relevant peer methods for missing value pre-imputation

- **Min/2 (half minimum):** Taking MNAR as the missing mechanism (e.g. LLOD), for each protein the missing values are estimated as half the minimum value of the observed intensities in that protein across all samples (Herrington, et al., 2018; Liu and Dongre, 2020).
- **Mean:** For MAR/MCAR as the missing values mechanism, for each protein we replaced the missing values with the mean value of the observed intensities in that protein across all samples (Herrington, et al., 2018; Liu and Dongre, 2020).

##### Design of simulation data and evaluation experiments

As we discussed in our ProImput paper (Shen, et al., 2022), for real omics data, there is no method that can truly assess the accuracy of various imputation methods, because missing values will never be known and masked values cannot serve as the ground-truth missing values for unbiased evaluation. To validate the efficacy of migImput, we evaluated the performance of imputations by migImput, Mean and Min/2 via well-controlled realistic simulation studies (Wu, et al., 2022). The simulations involve five groups, 1,000 genes, and 150 samples (30 samples for each group). Of the 1,000 genes, 60% (= 600) are 'asymmetric' differentially-expressed genes (aDEGs). The numbers of aDEGs among groups are set to be asymmetric (upregulated), e.g., 60, 90, 120, 150, 180 DEGs, distributed among the five groups, respectively. These aDEGs mimic both marker genes (MGs) and signature genes (SGs).

The ground-truth missing values were introduced and assigned by assigning some of the observed values with NA. The overall missing rate is 46.7%, that is, 70, 000 missing values out of total 150,000 entries. The overall missing rate due to LLOD is 6.7%, that is, 10,000 MNAR missing values. The overall missing rate due to MCAR is 40%, that is, 60,000 MCAR missing values. Theoretically, any gene may contain some missing values. Some of the simulated 1,000 genes with overall missing rates higher than a threshold (e.g. >60%) were 'masked out' to mimic 'marker or signature genes' that are often prematurely eliminated.

#### Results (with supplementary figures and tables)

##### migImput to recruit signature genes

**Table S1**. Imputation accuracy produced by migImput as compared with Mean and Min/2 via realistic simulation studies and measured by mean Euclidean distance.

| | Mean Euclidean distance |
|---|---|
| migImput | 17.9081 |
| Min/2 | 341.4434 |
| Mean | 324.0214 |

**Detection of eSGs by eCOT displayed via uniHM**

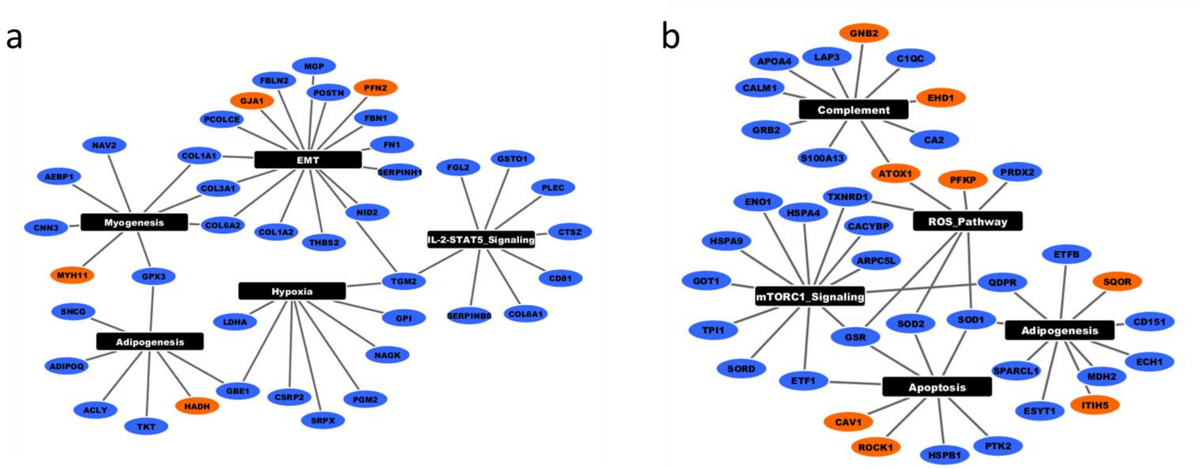

**Figure S1**. Upregulated (orange nodes) and downregulated (blue nodes) signature genes clustered into the top 5 functional pathways from the MSigDB component of Enrichr pathway analysis software (black nodes) are shown for the normal NL (A) and fatty streak FS (B) groups.

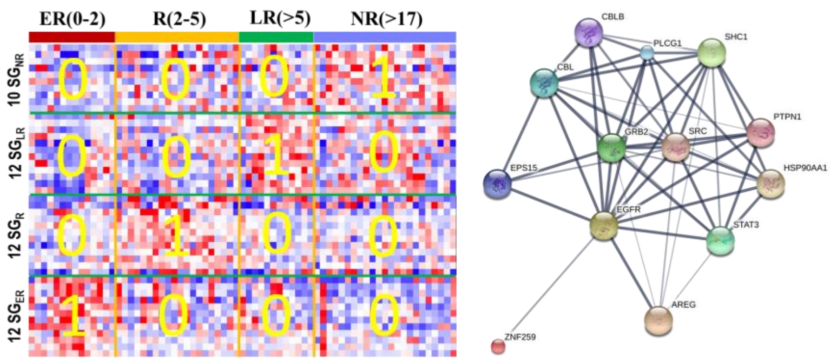

**Figure S2**. Signature genes associated with recurrence progression, detected by COT/eCOT and displayed by new heatmap design (Edinburgh breast cancer transcriptomics data), and a typical EGFR-ZNF259-AREG signaling network.

**Table S2**. The protein IDs of SGs associated with NL, FS, and FP, respectively, and their corresponding cosine scores readily provided by the COT/eCOT tool, on the proteomics dataset acquired from human artery samples enriched by the tissue types associated with atherosclerosis.

| Subtype-specific Signature Genes (SGs) | | | | | | | | | |
|---|---|---|---|---|---|---|---|---|---|
| $SG_{NL}$ | | $SG_{FS}$ | | $SG_{FP}$ | | | | | |
| SGs | Cosine Score | SGs | Cosine Score | SGs | Cosine Score | SGs | Cosine Score | SGs | Cosine Score |
| BANF1 | 0.66399901 | GNB2 | 0.79616713 | JCHAIN | 0.99706477 | C7 | 0.96885745 | CORO1A | 0.9478701 |
| PACSIN2 | 0.6279405 | NDRG1 | 0.73916782 | A2M | 0.99687182 | PGLYRP2 | 0.96851954 | C8A | 0.94614345 |
| PTPN11 | 0.62600027 | ITIH5 | 0.68218832 | PLTP | 0.99533245 | CTSB | 0.96843119 | CTSD | 0.94603303 |
| ACTN1 | 0.61876406 | EPB41L2 | 0.67824745 | HP | 0.99418856 | CPB2 | 0.9671609 | C3 | 0.94442917 |
| MYH11 | 0.61620071 | PSMD1 | 0.67263765 | SAA4 | 0.99265154 | IGHG1 | 0.96630768 | TYMP | 0.94438329 |
| TMEM43 | 0.61325619 | PSMA7 | 0.67116557 | FGG | 0.99192681 | C4BPB | 0.96597299 | C9 | 0.94424813 |
| GJA1 | 0.61182973 | NCKAP1 | 0.66207196 | IGHV3-23 | 0.99165202 | PLG | 0.96480473 | C4B | 0.94263471 |
| PFN2 | 0.60913077 | BCAM | 0.65589275 | FGB | 0.99117752 | F2 | 0.96361546 | AFM | 0.94240442 |
| HSD17B10 | 0.60870231 | CAV1 | 0.65569599 | IGKV3-11 | 0.99051544 | ITIH4 | 0.96272169 | C2 | 0.94151828 |
| MYL6 | 0.60805089 | DCTN1 | 0.65204032 | IGFALS | 0.98871268 | C8G | 0.96056103 | FABP5 | 0.94106945 |
| EML2 | 0.60792498 | SQOR | 0.64554379 | IGHA1 | 0.98555638 | CP | 0.959959 | APMAP | 0.93849627 |
| VCL | 0.60365104 | GDI1 | 0.64234243 | APOA1 | 0.9853748 | ITIH1 | 0.95937765 | F13A1 | 0.93806871 |
| HADH | 0.59959516 | CCT3 | 0.64108435 | APOE | 0.98509745 | C6 | 0.95899082 | C1R | 0.93740563 |
| LRRC47 | 0.59905334 | SNX2 | 0.64070421 | APOB | 0.98474853 | TNC | 0.95802925 | TF | 0.93718407 |
| | | ROCK1 | 0.63908302 | C4BPA | 0.98441775 | HRG | 0.95761387 | ORM1 | 0.93703461 |
| | | AOC3 | 0.63858726 | IGHM | 0.98401872 | SERPINA6 | 0.95702299 | C8B | 0.93680165 |
| | | MYADM | 0.63626786 | FGA | 0.98395106 | A1BG | 0.95608454 | THBS1 | 0.93626904 |
| | | LASP1 | 0.63617626 | IGHG4 | 0.98384512 | CLU | 0.95552078 | SERPING1 | 0.93536841 |
| | | ATP2B4 | 0.63571816 | ITIH2 | 0.98281986 | SERPIND1 | 0.9546262 | ECM1 | 0.93374806 |
| | | AP2A1 | 0.63563977 | IGLV3-19 | 0.98103357 | CTSG | 0.95328289 | LYZ | 0.93274026 |
| | | EHD1 | 0.63517103 | APOD | 0.98089455 | MB | 0.95324877 | HBD | 0.93249064 |
| | | SYNPO | 0.63381368 | VTN | 0.98071371 | C1S | 0.95296121 | SERPINC1 | 0.93043207 |
| | | SPTBN1 | 0.63330807 | IGKV4-1 | 0.98064623 | SERPINF2 | 0.95203776 | ALB | 0.92951055 |
| | | PFKP | 0.6331388 | IGKV2D-28 | 0.9779092 | KNG1 | 0.95195618 | AGT | 0.92901247 |
| | | DBNL | 0.63142294 | APOA2 | 0.97732094 | DAAM2 | 0.95185001 | HTRA1 | 0.92676544 |
| | | RTN4 | 0.63088827 | APOC3 | 0.97711365 | GC | 0.95183061 | SERPINA4 | 0.9265501 |
| | | PRKAR2A | 0.62799511 | QSOX1 | 0.97659695 | IGHV3-7 | 0.9514402 | PXDN | 0.92355079 |
| | | CORO1B | 0.62587838 | IGKV3-20 | 0.97517938 | TTR | 0.95020494 | MFAP5 | 0.92268903 |
| | | CAPNS1 | 0.62579346 | IGHV3-53 | 0.9750816 | C5 | 0.95007188 | SERPINA3 | 0.91774302 |
| | | SORBS3 | 0.62557177 | IGKC | 0.97453607 | APCS | 0.94992204 | F9 | 0.91533295 |
| | | DYNC1H1 | 0.62538578 | IGHG3 | 0.97235091 | CFI | 0.94840171 | IGHG2 | 0.91417815 |
| | | ATOX1 | 0.62416812 | IGLV3-21 | 0.97024749 | CFH | 0.94810627 | LRG1 | 0.91297612 |
| | | VAT1 | 0.62408469 | IGKV1-39 | 0.96887079 | SERPINA1 | 0.94796173 | AHSG | 0.91243482 |

**Table S3**. The protein IDs of DSGs associated with NL, FS, and FP, respectively, and their corresponding cosine scores readily provided by the COT/eCOT tool, on the proteomics dataset acquired from human artery samples enriched by the tissue types associated with atherosclerosis.

| Down-regulated Subtype-specific Signature Genes (DSGs) | | | | | | | | | | | |
|---|---|---|---|---|---|---|---|---|---|---|---|
| DSGNL | | | | DSGFS | | | | DSGFP | | | |
| DSGs | Cosine Score | DSGs | Cosine Score | DSGs | Cosine Score | DSGs | Cosine Score | DSGs | Cosine Score | DSGs | Cosine Score |
| UGDH | 0.97796828 | GPNMB | 0.91328912 | RPL11 | 0.96298743 | HNRNPC | 0.87185927 | SYNM | 0.98871309 | PALLD | 0.88236194 |
| POSTN | 0.97790452 | EMD | 0.91320961 | DNAJC10 | 0.95490725 | TPI1 | 0.87159977 | DES | 0.96044156 | CNN1 | 0.87925931 |
| ACLY | 0.97458651 | RPL18A | 0.91319175 | LOXL1 | 0.92561811 | EEF1D | 0.87097148 | VASP | 0.96026919 | PDLIM4 | 0.87791023 |
| SNCG | 0.97163721 | ARPC1B | 0.91303968 | MYL9 | 0.9198815 | HNRNPD | 0.87064768 | DDX1 | 0.95771017 | OXCT1 | 0.87764782 |
| FRZB | 0.9640618 | CFHR2 | 0.91292782 | DDOST | 0.91646524 | RSU1 | 0.87063661 | CSPG4 | 0.95684982 | PDLIM5 | 0.87444761 |
| CSRP2 | 0.94768822 | AKR7A2 | 0.91290705 | LIMS1 | 0.90817943 | TXNDC5 | 0.87054949 | PDHB | 0.95681944 | PRKG1 | 0.87401774 |
| B2M | 0.94626952 | FBLN2 | 0.91274726 | FBLN5 | 0.90656249 | P4HB | 0.86948238 | TLN2 | 0.94996674 | FHL1 | 0.87326865 |
| SAMHD1 | 0.9433037 | PKM | 0.91269418 | VCAN | 0.90600331 | CSTB | 0.86945827 | MAP4 | 0.94583419 | LPP | 0.87287757 |
| SERPINB6 | 0.94325709 | TPP1 | 0.91208321 | MDH2 | 0.90522063 | HSPA8 | 0.8692552 | PFKM | 0.94046164 | CSRP1 | 0.87269445 |
| FLNB | 0.94143511 | TAGLN2 | 0.91183794 | ADGRE5 | 0.90340783 | ACTA1 | 0.86907756 | PCBD1 | 0.93735518 | SCRN1 | 0.87188634 |
| HSP90AB1 | 0.94121487 | LDHA | 0.91180937 | PSAP | 0.89945747 | ARPC3 | 0.86892634 | MAP1B | 0.9363296 | CORO1C | 0.87129673 |
| ADH1B | 0.93943507 | SERPINH1 | 0.91156582 | HSPA4 | 0.89883762 | RPLP1 | 0.86880872 | SMTN | 0.93616209 | LMCD1 | 0.87051231 |
| FHL2 | 0.93829958 | SRPX | 0.91135931 | SOD3 | 0.89857818 | APOA4 | 0.86866414 | SYNPO2 | 0.93585612 | ZYX | 0.8692396 |
| THBS2 | 0.93683007 | HSP90B1 | 0.91109945 | CALM1 | 0.89739642 | H2AC21 | 0.86770686 | KRT8 | 0.93471255 | MAOB | 0.86899583 |
| GBE1 | 0.93658625 | PGM2 | 0.91083891 | LGALS3BP | 0.89733833 | GUK1 | 0.8676276 | GPD1L | 0.93381032 | SORBS2 | 0.8685885 |
| CNN3 | 0.93535819 | COL6A2 | 0.91031765 | TXNRD1 | 0.89717995 | SOD1 | 0.86742788 | PDLIM1 | 0.93166183 | LMOD1 | 0.86852852 |
| RPS5 | 0.93493486 | ARHGDIB | 0.91025203 | ELAVL1 | 0.89397408 | LGALS1 | 0.8672633 | SLMAP | 0.9313632 | PGM5 | 0.86642102 |
| ASPN | 0.93269194 | AKR1B1 | 0.91007335 | MFAP4 | 0.89332732 | SPARCL1 | 0.8669548 | KANK2 | 0.93024388 | CAST | 0.86597785 |
| RPS3A | 0.93181656 | POTEI | 0.91004027 | MPST | 0.89186194 | ATIC | 0.86682776 | ALDH1B1 | 0.92654204 | PFKL | 0.86577581 |
| FGL2 | 0.92994064 | RPL10A | 0.90995312 | LAP3 | 0.89175485 | COTL1 | 0.86658285 | ATL3 | 0.92194334 | PDLIM3 | 0.86421523 |
| FN1 | 0.92984785 | DHX9 | 0.90948583 | PTK2 | 0.89139498 | EEF1G | 0.86597648 | NEXN | 0.92164605 | ACAT1 | 0.86263986 |
| CTSZ | 0.92967462 | CPA3 | 0.9093765 | PDCD6 | 0.89111401 | HBB | 0.86578032 | TMOD1 | 0.91889264 | FLNA | 0.86099051 |
| ACTBL2 | 0.92804321 | POTEF | 0.90923204 | GOT1 | 0.88979012 | MAPK1 | 0.86528056 | CPNE3 | 0.91700374 | COL18A1 | 0.86056826 |
| COL1A1 | 0.9276712 | PSMA2 | 0.90905685 | GRB2 | 0.88967176 | GSR | 0.8650212 | CKB | 0.91616665 | TGFB1I1 | 0.8598698 |
| CDC42 | 0.92755506 | SNX6 | 0.90842419 | SORD | 0.88958989 | PEBP1 | 0.86459692 | UBR4 | 0.91496578 | PARVA | 0.85936976 |
| COL14A1 | 0.92718802 | GPX3 | 0.9076328 | ARPC5L | 0.88946186 | S100A13 | 0.86451566 | FLNC | 0.9133149 | NUDC | 0.85915921 |
| GPI | 0.92515466 | AEBP1 | 0.90744514 | SLC4A1 | 0.88823332 | AGL | 0.86441902 | DMD | 0.91048156 | CAP2 | 0.85910847 |
| ORM2 | 0.92301675 | CISD1 | 0.90737094 | HSPA9 | 0.88650987 | MT-CO2 | 0.86390956 | MAPRE1 | 0.90354419 | PRKRA | 0.8585306 |
| TNXB | 0.92296932 | CRYZ | 0.90686408 | KIF5B | 0.8853924 | HSPB1 | 0.8638633 | ITGA7 | 0.90283697 | DARS1 | 0.85794006 |
| LAMB1 | 0.92247931 | COL3A1 | 0.90684804 | ETFB | 0.8848053 | RRAS | 0.8630933 | SORBS1 | 0.90183205 | OLA1 | 0.85786658 |
| COL1A2 | 0.92195768 | COX4I1 | 0.9067562 | LAMP1 | 0.88353912 | ITGB5 | 0.86308701 | HSPB6 | 0.90033356 | MYLK | 0.85457049 |
| CA3 | 0.92052789 | F12 | 0.90578917 | H1-10 | 0.88353436 | PRDX2 | 0.86298843 | CALD1 | 0.90014832 | DSTN | 0.85401586 |
| NAV2 | 0.91972431 | PURA | 0.90554382 | NDUFA2 | 0.88332684 | MAT2B | 0.86295257 | ACAN | 0.90003168 | PLIN3 | 0.85255948 |
| TGM2 | 0.91960481 | BLVRA | 0.90532886 | H4C1 | 0.88068154 | LTBP4 | 0.86282416 | KTN1 | 0.89965604 | IPO7 | 0.85217264 |
| RPS6 | 0.9186658 | TKT | 0.9052027 | RAB10 | 0.87974357 | COL21A1 | 0.86250159 | EFHD1 | 0.89801688 | RBPMS | 0.85194024 |
| PCOLCE | 0.91835886 | COL6A1 | 0.90482364 | PPP1CB | 0.87930131 | CACYBP | 0.86238299 | SGCD | 0.89714648 | PTGIS | 0.85175904 |
| BCAP31 | 0.91823109 | HP1BP3 | 0.904794 | CA2 | 0.87862213 | ESYT1 | 0.86229239 | MCAM | 0.89593728 | ARPC1A | 0.85162258 |
| NID2 | 0.91806208 | CPXM2 | 0.90449191 | CFHR1 | 0.87815208 | ERP44 | 0.86192676 | TUBB6 | 0.8940541 | HAAO | 0.85150942 |
| MGP | 0.91759981 | PLXDC2 | 0.90412973 | SOD2 | 0.87792655 | SUN2 | 0.86168264 | TPM2 | 0.89329357 | TES | 0.85131881 |
| ARPC2 | 0.91739988 | PDIA4 | 0.90384272 | HBA1 | 0.87777999 | CZIB | 0.86146952 | UQCRC2 | 0.89090684 | ITGA8 | 0.85098183 |
| CYBRD1 | 0.91690673 | PLEC | 0.90372038 | KRT10 | 0.87725099 | C1QC | 0.86110193 | SCUBE3 | 0.88803634 | SND1 | 0.85049133 |
| FBN1 | 0.91639543 | CD81 | 0.90363896 | CCN3 | 0.87580467 | ARHGDIA | 0.86090797 | TNS1 | 0.88776552 | TPM1 | 0.85032875 |
| ADIPOQ | 0.91486101 | NAGK | 0.90354618 | CD151 | 0.87506969 | CRYAB | 0.8606558 | SNTB2 | 0.8870108 | KPNB1 | 0.85012221 |
| SYPL1 | 0.91480787 | COPB2 | 0.90315964 | ETF1 | 0.87462863 | PODN | 0.86059141 | LDB3 | 0.88647079 | PRKAR1A | 0.84862744 |
| GSTO1 | 0.91479581 | RPS9 | 0.90313816 | ENO1 | 0.8741981 | OGN | 0.86058936 | TBCA | 0.88646615 | PEA15 | 0.84765444 |
| SEPTIN11 | 0.91479243 | TINAGL1 | 0.9029865 | ELOB | 0.8736641 | SELENBP1 | 0.86048369 | CLIC4 | 0.88646589 | VIM | 0.84737145 |
| CLEC3B | 0.91429825 | CRKL | 0.90266835 | PSMA6 | 0.8732705 | QDPR | 0.8602868 | LMNB1 | 0.88625535 | TAGLN | 0.84607289 |
| RPSA | 0.91419453 | RACK1 | 0.90259484 | SELENOM | 0.87295461 | ATP6V1A | 0.86009563 | SPON1 | 0.88497963 | TPD52L2 | 0.84598021 |
| VWF | 0.91369272 | TNFSF13 | 0.90245735 | ECH1 | 0.87256888 | PSMB6 | 0.85990107 | PDLIM7 | 0.88443033 | VDAC2 | 0.84568603 |
| ARF6 | 0.91349677 | | | LTBP2 | 0.87203069 | RPL7 | 0.85988186 | IMMT | 0.88329922 | | |

| | | |
|---|---|---|
| Total Patients | : | 110 |
| No. of microarray samples | : | 110 |
| Sample Collection | : | Pretreatment |

| Edinburg First Batch Samples | | | |
|---|---|---|---|
| Recurred (46) | (0-2) years | (2–5) years | >5 years |
| | 14 | 20 | 12 |
| Non Recurred (NR) (64) | (5.9-11) years | (11–17) years | >17 years |
| | Not Considered | | 23 |

| Reference | Number | Quality |
|---|---|---|
| 1 1 1 0 | 30 | 0.89 |
| 1 1 0 1 | 30 | 0.89 |
| 1 0 1 1 | 30 | 0.89 |
| 0 1 1 1 | 30 | 0.90 |

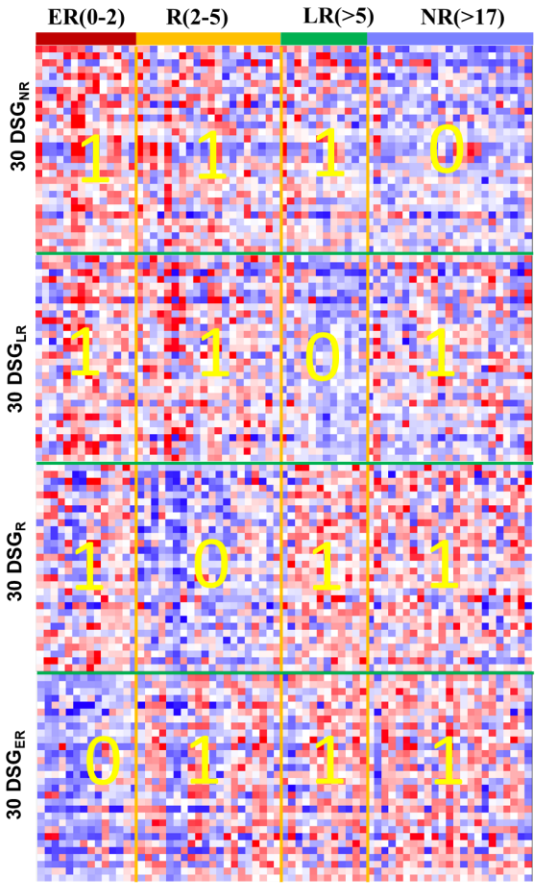

**Figure S3**. The DSGs detected by COT/eCOT on the Edinburgh breast cancer gene expression data that were acquired prior to standard treatment. The DSGs are displayed via new heatmap design and associated with ER (early recurrence, 0~2 years), R (recurrence, 2~5 years), LR (late recurrence, >5 years), and NR (never recurrence, >17 years), respectively.

| Reference | Number | Quality |
|---|---|---|
| 0 0 1 1 | 15 | 0.77 |
| 1 1 0 0 | 15 | 0.76 |
| 0 1 1 0 | 10 | 0.76 |
| 1 0 0 1 | 10 | 0.75 |
| 1 0 1 0 | 10 | 0.76 |
| 0 1 0 1 | 10 | 0.76 |

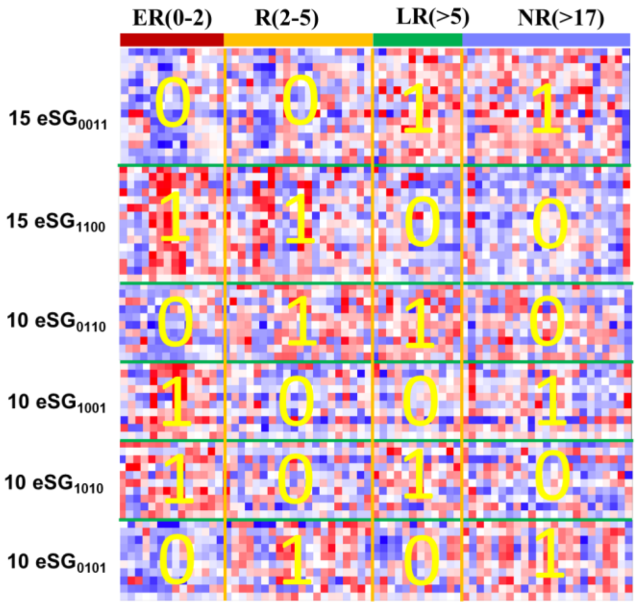

**Figure S4**. The CSGs detected by eCOT on the Edinburgh breast cancer gene expression data that were acquired prior to standard treatment. The eSGs are displayed via new heatmap design and associated with ER, R, LR, and NR, respectively.

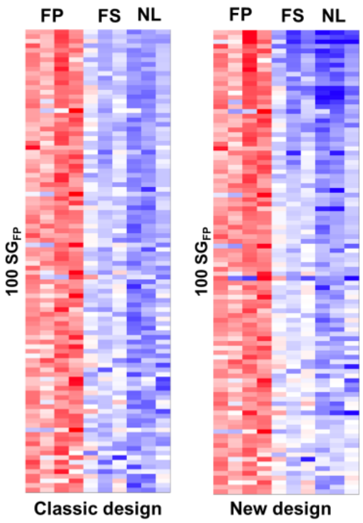

**Figure S5.** A side-by-side display of comparing the classic heatmap and the new heatmap, on the 100 protein SGs associated with FP.

**Table S4**. The SGs associated with ER, R, LR, and NR, respectively, and their corresponding cosine scores readily provided by the COT/eCOT tool, on the Edinburgh breast cancer gene expression data that were acquired prior to standard treatment.

| 10 $SG_{NR}$ | | 12 $SG_{LR}$ | | 12 $SG_{R}$ | | 12 $SG_{ER}$ | |
|---|---|---|---|---|---|---|---|
| Gene names | Cosine Score | Gene names | Cosine Score | Gene names | Cosine Score | Gene names | Cosine Score |
| --- | 0.57 | --- | 0.58 | SYT1 | 0.58 | --- | 0.61 |
| NPY1R | 0.57 | --- | 0.58 | --- | 0.56 | CP | 0.60 |
| NOVA1 | 0.56 | RBM24 | 0.58 | LOC100129518 | 0.56 | CP | 0.60 |
| CPB1 | 0.55 | LINC01410 | 0.58 | ERN1 | 0.55 | SOX11 | 0.60 |
| GLYATL1 | 0.55 | FLJ38379 | 0.57 | RNA45S5 | 0.55 | GPR126 | 0.59 |
| LOC101926959 | 0.55 | MUM1L1 | 0.57 | DNHD1 | 0.55 | PSAT1 | 0.59 |
| NEFH | 0.55 | AREG | 0.57 | RP11-190A12.8 | 0.55 | S100A8 | 0.58 |
| ST6GALNAC5 | 0.55 | ELOVL2 | 0.57 | NEGR1 | 0.55 | CECR2 | 0.58 |
| --- | 0.55 | ABCC13 | 0.57 | LSS | 0.55 | DSC3 | 0.58 |
| GP2 | 0.55 | PTHLH | 0.57 | ERAP2 | 0.55 | CDCA7 | 0.57 |
| | | PGR | 0.57 | ELL2 | 0.55 | CHRM3 | 0.57 |
| | | CLSTN2 | 0.57 | BMPR1B | 0.55 | PLCH1 | 0.57 |

**Table S5**. The DSGs associated with ER, R, LR, and NR, respectively, and their corresponding cosine scores readily provided by the COT/eCOT tool, on the Edinburgh breast cancer gene expression data that were acquired prior to standard treatment.

| 30 $DSG_{NR}$ | | 30 $DSG_{LR}$ | | 30 $DSG_{R}$ | | 30 $DSG_{ER}$ | |
|---|---|---|---|---|---|---|---|
| Gene names | Cosine Score | Gene names | Cosine Score | Gene names | Cosine Score | Gene names | Cosine Score |
| CECR2 | 0.91 | PLEKHB1 | 0.90 | ABCC13 | 0.90 | --- | 0.92 |
| CP | 0.91 | HLA-DRB4 | 0.90 | LRP2 | 0.90 | GATA3-AS1 | 0.92 |
| DSCAM-AS1 | 0.90 | CXCL13 | 0.90 | GABRP | 0.90 | --- | 0.92 |
| S100P | 0.90 | SCNN1G | 0.90 | LINC00472 | 0.90 | GLRB | 0.91 |
| CP | 0.90 | CTSC | 0.90 | LINC01410 | 0.90 | NPY1R | 0.91 |
| PCOLCE2 | 0.90 | ADAMDEC1 | 0.90 | PROSER2 | 0.90 | FLJ38379 | 0.91 |
| GPR126 | 0.90 | FCRL5 | 0.90 | RASSF10 | 0.90 | PTGER3 | 0.91 |
| ZNF595 | 0.90 | MIR6787 | 0.90 | FBN2 | 0.90 | BMPR1B | 0.91 |
| SOX11 | 0.89 | UNC5A | 0.90 | PCDHA1 | 0.90 | NEFH | 0.91 |
| THRSP | 0.89 | SULT1C2 | 0.89 | LINC00472 | 0.90 | GAD1 | 0.91 |
| S100A8 | 0.89 | RARRES1 | 0.89 | MAOB | 0.90 | KCNJ3 | 0.91 |
| TDO2 | 0.89 | ZNF204P | 0.89 | KRT23 | 0.90 | PGR | 0.91 |
| CPT1A | 0.89 | RARRES1 | 0.89 | --- | 0.90 | FLJ38379 | 0.91 |
| CP | 0.89 | LOC100129518 | 0.89 | LOC101926959 | 0.89 | --- | 0.91 |
| CLCA2 | 0.89 | HIST1H2BC | 0.89 | SHC4 | 0.89 | TTC6 | 0.91 |
| CLCA2 | 0.89 | CCAR1 | 0.89 | --- | 0.89 | MUM1L1 | 0.91 |
| MUCL1 | 0.89 | SEMA6A | 0.89 | WNK4 | 0.89 | BMPR1B | 0.91 |
| CELF5 | 0.89 | FDCSP | 0.89 | SLC38A1 | 0.89 | AGTR1 | 0.90 |
| SOX11 | 0.89 | ARG2 | 0.89 | TMEM139 | 0.89 | PTGER3 | 0.90 |
| BEND7 | 0.89 | RPUSD3 | 0.89 | NDP | 0.89 | HLA-DQB1 | 0.90 |
| GSDMB | 0.89 | --- | 0.89 | TMEM45B | 0.89 | LINC01116 | 0.90 |
| --- | 0.89 | C6orf141 | 0.89 | GRP | 0.89 | SYT1 | 0.90 |
| COL4A3 | 0.89 | --- | 0.89 | PHOSPHO2 | 0.89 | PCAT18 | 0.90 |
| GLYATL2 | 0.89 | BCL11A | 0.89 | ID4 | 0.89 | CPB1 | 0.90 |
| HOTAIR | 0.89 | HOXB6 | 0.89 | LRP2 | 0.89 | SERPINA1 | 0.90 |
| FAR2 | 0.89 | SLC2A5 | 0.89 | PRKAR2B | 0.89 | HLA-DQA1 | 0.90 |
| KLHDC7B | 0.89 | SOX11 | 0.89 | TINCR | 0.89 | ADCY1 | 0.90 |
| GPRC5A | 0.89 | BCL11A | 0.89 | FRZB | 0.89 | HLA-DQA1 | 0.90 |
| --- | 0.89 | ST8SIA1 | 0.89 | ARX | 0.89 | LOC692247 | 0.90 |
| STC1 | 0.89 | PLA2G7 | 0.89 | TM4SF18 | 0.89 | GREB1L | 0.90 |

**Table S6**. The eSGs associated with ER, R, LR, and NR, respectively, and their corresponding cosine scores readily provided by the eCOT tool, on the Edinburgh breast cancer gene expression data that were acquired prior to standard treatment.

| 15 $eSG_{0011}$ | | 15 $eSG_{1100}$ | | 15 $eSG_{0110}$ | | 10 $eSG_{1001}$ | | 10 $eSG_{1010}$ | | 10 $eSG_{0101}$ | |
|---|---|---|---|---|---|---|---|---|---|---|---|
| Gene names | Cosine Score | Gene names | Cosine Score | Gene names | Cosine Score | Gene names | Cosine Score | Gene names | Cosine Score | Gene names | Cosine Score |
| PGR | 0.78 | SOX11 | 0.79 | SYT1 | 0.77 | PSAT1 | 0.75 | CECR2 | 0.77 | NPY1R | 0.78 |
| NDP | 0.78 | CP | 0.78 | ADIPOQ | 0.76 | PROM1 | 0.75 | ABCC13 | 0.77 | GP2 | 0.77 |
| ENPP5 | 0.77 | FCRL5 | 0.78 | DSCAM-AS1 | 0.76 | DSC3 | 0.75 | --- | 0.77 | BMPR1B | 0.76 |
| LINC00472 | 0.77 | S100A8 | 0.77 | ADH1B | 0.76 | CHRM3 | 0.75 | ARX | 0.77 | --- | 0.76 |
| GLRB | 0.77 | LOC100129518 | 0.77 | GATA3-AS1 | 0.76 | GABRP | 0.75 | TMEM45B | 0.76 | BMPR1B | 0.76 |
| PTHLH | 0.77 | SULT1C2 | 0.77 | FLJ38379 | 0.76 | FDCSP | 0.75 | LINC01410 | 0.76 | NOVA1 | 0.76 |
| CLIC6 | 0.77 | CP | 0.77 | PCAT18 | 0.76 | ACN9 | 0.75 | CP | 0.76 | NEGR1 | 0.76 |
| --- | 0.77 | RARRES1 | 0.76 | KCNJ3 | 0.76 | --- | 0.75 | GPR126 | 0.76 | AGTR1 | 0.75 |
| CPB1 | 0.77 | --- | 0.76 | CTD-2311B13.7 | 0.76 | TTC22 | 0.75 | HOTAIR | 0.76 | HLA-DQA1 | 0.75 |
| CLSTN2 | 0.77 | SLAMF7 | 0.76 | --- | 0.75 | HEY2 | 0.75 | TMSB15A | 0.76 | NCR3LG1 | 0.75 |
| RBM24 | 0.77 | CXCL13 | 0.76 | | | | | | | | |
| --- | 0.77 | ST8SIA1 | 0.76 | | | | | | | | |
| AFF3 | 0.77 | BCL11A | 0.76 | | | | | | | | |
| PTGER3 | 0.77 | GPR126 | 0.76 | | | | | | | | |
| --- | 0.77 | RNA45S5 | 0.76 | | | | | | | | |

### Discussion

The ability to simulate the missing values mechanisms (MNAR, MAR/MCAR) depends on the efficacy of the tools applied. While it may be informative to compare the impact of the imputation versus non-imputation on some subsequent data analysis, we have opted to focus on assessing direct imputation accuracy using realistic simulations with available ground truth, because the evaluation using subsequent analysis would be indirect and task-dependent. In future work, support vector machine or artificial neural network (ANN) based methods may be considered as emerging imputation competitors (Wang, et al., 2020), and a combination approach utilizing an ensemble of strategies could be explored (Ma, et al., 2020). We have recently begun to explore a deep matrix completion method (Fan, et al., 2021).

The null distribution plays a crucial role in large-scale multiple testing when false positives are of great concern. However, because the number of pure group samples is often very small and eSG/non-eSG patterns are often highly complex and intrinsically data-dependent, classical schemes to estimate the null distribution in a two/multiple-sample test setting is impractical (Chen, et al., 2021) or even inappropriate (Efron, 2004). A reasonable assumption is that the observed data can show the null distribution when a significant majority of features are associated with the null hypothesis (Efron, 2004).

Functional pathway analysis of all signature genes, both upregulated (SG) and down-regulated (DSG), performed on each pathological group produced results consistent with known pathogenesis in atherogenesis. Network analysis of the top enriched functional pathways from analysis of fibrous plaque genes indicated upregulated SGs were enriched for complement and coagulation functions, whereas DSGs were enriched for Myogenesis and EMT (**Fig. 3**). Together, this pattern is consistent with increased inflammation and decreased smooth muscle cell contractile phenotype composition within atherosclerotic lesions (PMID: 32202800, PMID: 35365353, PMC4762053, PMID: 15269336). While equal numbers of SG and DSGs were confidently identified for the FP group, there was lower SG quality (e.g., lower cosine score) and fewer numbers identified for the FS and NL groups, although DSG for these groups were strong. Pathway analysis for the FS and NL groups indicated marker genes associated with mTORC1 signaling and reactive oxygen species (ROS) pathway enriched among FS signature proteins and myogenesis, EMT, hypoxia and IL2/STAT5 signaling were enriched among NL signature proteins (**Fig. S1**). Both mTORC1 and ROShave previously been linked to atherogenesis (PMC8835022, PMID:

28446473). Interestingly, IL2 in blood vessels is produced, at least in part, by resident T cells with IL2 receptors located on the smooth muscle cells (PMC3162067) and IL2 signaling has been linked to atherogenesis (PMID: 1515626). Since IL2-related proteins were enriched among the DSG in the NL group, this finding could reflect that lower IL2 signaling is protective against atherosclerotic plaque development. This hypothesis and others generated from the analysis of SG and DSG warrants further testing in future studies.

#### R Scripts

The R scripts are available at https://github.com/niccolodpdu/ABDS

More suggestions on parameter setting can be found in the package vignette.

### References

Chen, L., *et al.* Data-driven detection of subtype-specific differentially expressed genes. *Scientific Reports* 2021;11:332.

Chen, L., *et al.* debCAM: a bioconductor R package for fully unsupervised deconvolution of complex tissues. *Bioinformatics* 2020;36(12):3927-3929.

Chikina, M., Zaslavsky, E. and Sealfon, S.C. CellCODE: a robust latent variable approach to differential expression analysis for heterogeneous cell populations. *Bioinformatics* 2015;31(10):1584-1591.

Clarke, R., *et al.* The properties of high-dimensional data spaces: implications for exploring gene and protein expression data. *Nat Rev Cancer* 2008;8(1):37-49.

Dabke, K., *et al.* A Simple Optimization Workflow to Enable Precise and Accurate Imputation of Missing Values in Proteomic Data Sets. *J. Proteome Res.* 2021;20(6):3214-3229.

Delaney, C., *et al.* Combinatorial prediction of marker panels from single-cell transcriptomic data. *Mol Syst Biol* 2019;15(10):e9005.

Efron, B. Large-scale simultaneous hypothesis testing: the choice of a null hypothesis. *Journal of the American Statistical Association* 2004;99(465):96-104.

Fan, M., *et al.* A deep matrix completion method for imputing missing histological data in breast cancer by integrating DCE-MRI radiomics. *Med Phys* 2021.

Herrington, D.M., *et al.* Proteomic Architecture of Human Coronary and Aortic Atherosclerosis. *Circulation* 2018;137(25):2741-2756.

Jakobsen, J.C., *et al.* When and how should multiple imputation be used for handling missing data in randomised clinical trials - a practical guide with flowcharts. *BMC Med Res Methodol* 2017;17(1):162.

Kuhn, A., *et al.* Population-specific expression analysis (PSEA) reveals molecular changes in diseased brain. *Nat Methods* 2011;8(11):945-947.

Lazar, C., *et al.* Accounting for the multiple natures of missing values in label-free quantitative proteomics data sets to compare imputation strategies. *Journal of proteome research* 2016;15(4):1116-1125.

Lin, X. and Boutros, P.C. Optimization and expansion of non-negative matrix factorization. *BMC Bioinformatics* 2020;21(1):7.

Liu, M. and Dongre, A. Proper imputation of missing values in proteomics datasets for differential expression analysis. *Brief Bioinform* 2020.

Lu, Y., *et al.* COT: an efficient and accurate method for detecting marker genes among many subtypes. *Bioinform Adv* 2022;2(1):vbac037.

Ma, H., *et al.* Recommender systems with social regularization. In, *The fourth ACM international conference on Web search and data mining*. Hong Kong: ACM Press; 2011. p. 287-296.

Ma, W., al., e. and Wang, P. DreamAI: algorithm for the imputation of proteomics data. *bioRxiv* 2020.

Newman, A.M., *et al.* Robust enumeration of cell subsets from tissue expression profiles. *Nat Methods* 2015;12(5):453-457.

Oba, S., *et al.* A Bayesian missing value estimation method for gene expression profile data. *Bioinformatics* 2003;19(16):2088-2096.

Parker, S.J., *et al.* Identification of putative early atherosclerosis biomarkers by unsupervised deconvolution of heterogeneous vascular proteomes. *J Proteome Res* 2020;19(7):2794-2806.

Ritchie, M.E., *et al.* limma powers differential expression analyses for RNA-sequencing and microarray studies. *Nucleic Acids Res* 2015;43(7):e47.
Shen, M., *et al.* Comparative assessment and novel strategy on methods for imputing proteomics data. *Sci Rep* 2022;12(1):1067.
Stekhoven, D.J. and Bühlmann, P. MissForest—non-parametric missing value imputation for mixed-type data. *Bioinformatics* 2012;28(1):112-118.
Wang, N., *et al.* Mathematical modelling of transcriptional heterogeneity identifies novel markers and subpopulations in complex tissues. *Scientific Reports* 2016;6:18909.
Wang, S., *et al.* NAguideR: performing and prioritizing missing value imputations for consistent bottom-up proteomic analyses. *Nucleic Acids Res* 2020;48(14):e83.
Webb-Robertson, B.-J.M., *et al.* Review, evaluation, and discussion of the challenges of missing value imputation for mass spectrometry-based label-free global proteomics. *Journal of proteome research* 2015;14(5):1993-2001.
Wu, C.-T., *et al.* Cosbin: Cosine score based iterative normalization of biologically diverse samples. *Bioinformatics Adv* 2022;3:vbac076.
